## Supplementary material for "Ventral pallidum GABA and glutamate neurons drive approach and avoidance through distinct modulation of VTA cell types": Figures S1 to S7 & Tables S1 to S2

Lauren Faget *et al.*

**This PDF file includes:**

Figs. S1 to S7
Tables S1 to S2

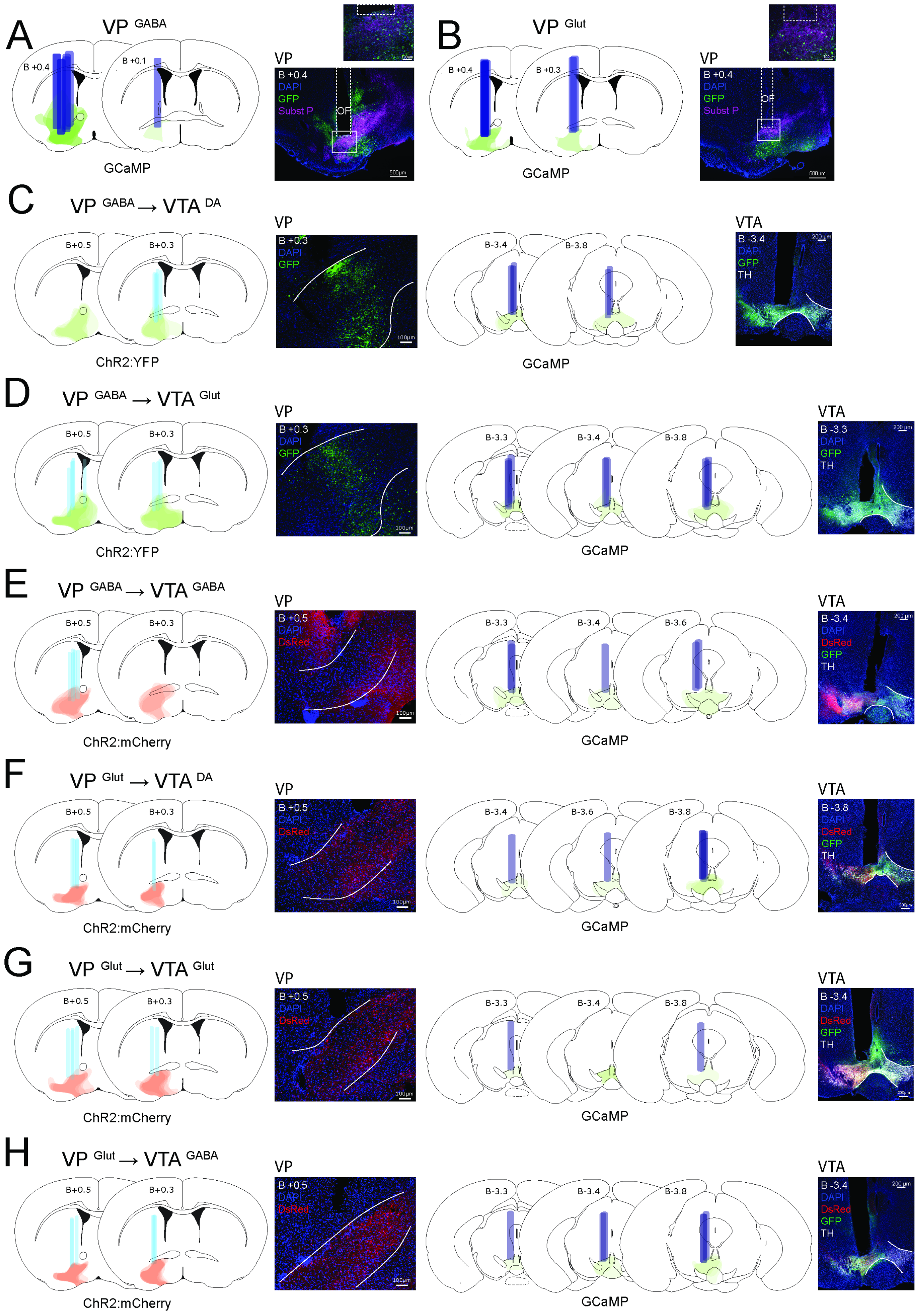

A
VP GABA
B +0.4
B +0.1
GCaMP
VP
B +0.4
DAPI
GFP
Subst P
OF
B
VP Glut
B +0.4
B +0.3
GCaMP
C
VP GABA → VTA DA
B+0.5
B+0.3
ChR2:YFP
B-3.4
B-3.8
GCaMP
VTA
DAPI
GFP
TH
D
VP GABA → VTA Glut
B-3.3
B-3.4
B-3.8
E
VP GABA → VTA GABA
ChR2:mCherry
DsRed
B-3.6
F
VP Glut → VTA DA
G
VP Glut → VTA Glut
H
VP Glut → VTA GABA

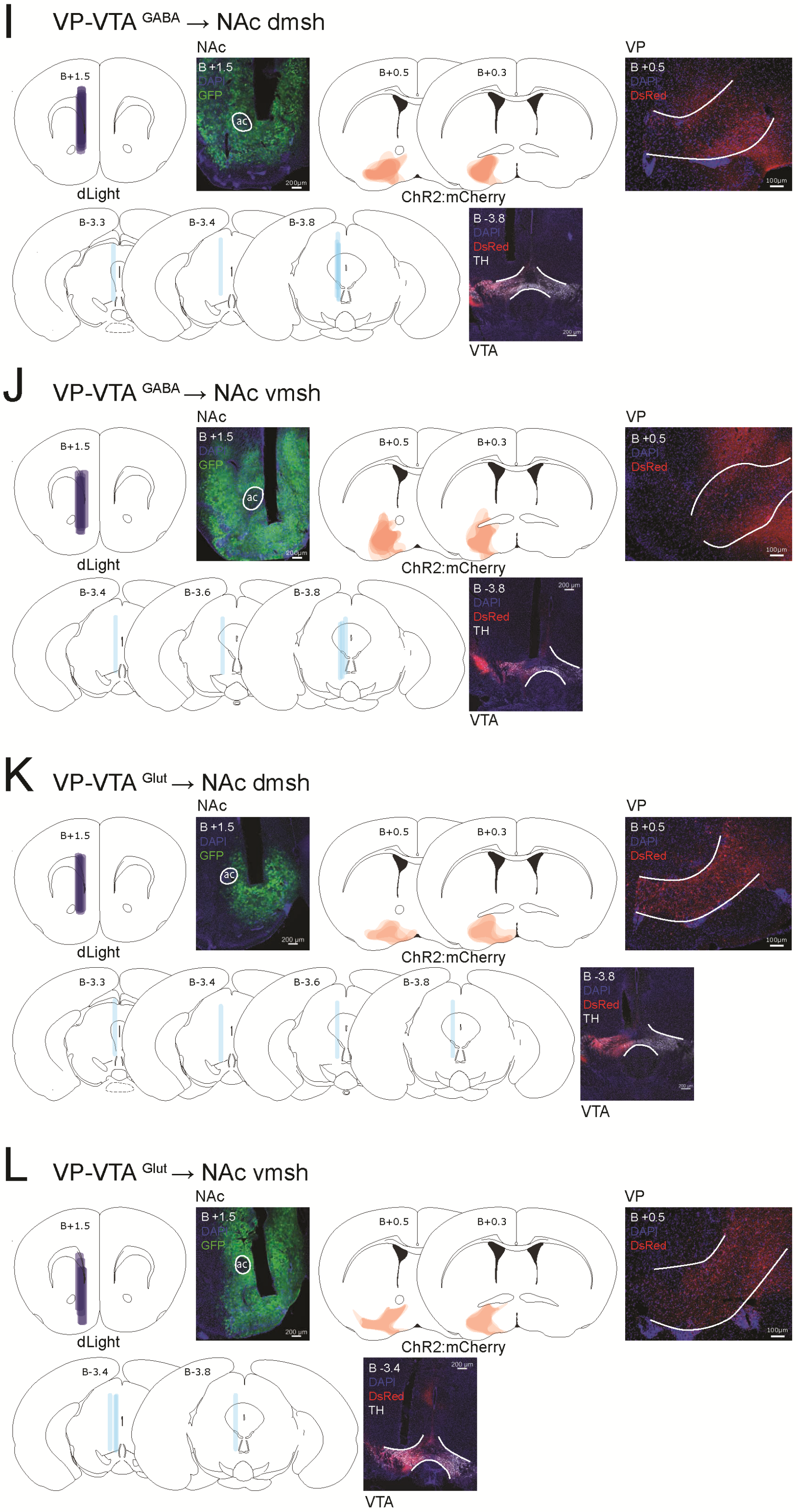
I
VP-VTA GABA → NAc dmsh
NAc
B+1.5
B +1.5
DAPI
GFP
ac
200μm
B+0.5
B+0.3
VP
B +0.5
DAPI
DsRed
100μm
dLight
ChR2:mCherry
B-3.3
B-3.4
B-3.8
B -3.8
DAPI
DsRed
TH
200 μm
VTA
J
VP-VTA GABA → NAc vmsh
NAc
B+1.5
B +1.5
DAPI
GFP
ac
200μm
B+0.5
B+0.3
VP
B +0.5
DAPI
DsRed
100μm
dLight
ChR2:mCherry
B-3.4
B-3.6
B-3.8
B -3.8
DAPI
DsRed
TH
200 μm
VTA
K
VP-VTA Glut → NAc dmsh
NAc
B+1.5
B +1.5
DAPI
GFP
ac
200 μm
B+0.5
B+0.3
VP
B +0.5
DAPI
DsRed
100μm
dLight
ChR2:mCherry
B-3.3
B-3.4
B-3.6
B-3.8
B -3.8
DAPI
DsRed
TH
200 μm
VTA
L
VP-VTA Glut → NAc vmsh
NAc
B+1.5
B +1.5
DAPI
GFP
ac
200 μm
B+0.5
B+0.3
VP
B +0.5
DAPI
DsRed
100μm
dLight
ChR2:mCherry
B-3.4
B-3.8
B -3.4
DAPI
DsRed
TH
200 μm
VTA

**Fig. S1. Histological validation of reporter/opsin expression and fiber tract placements. A-L.** Coronal sections showing example images of GCaMP reporter in VP or VTA, ChR2 opsin in VP or VTA, and/or dLight reporter in NAc for each experiment as indicated. GFP and DsRed immunoreactivity amplified viral GFP/YFP and mCherry fluorescence respectively. Some images show substance P immunoreactivity used to demark VP borders, Tyrosine hydroxylase (TH) to demark VTA, and DAPI nuclear stain is shown in blue. Diagrams schematize spread of indicated opsin or reporter and optic fiber (OF) placements at various points relative to Bregma (B) in mm; ac, anterior commissure. See Table S1 for details on viral and genetic strategies. Related to Fig. 1 to 6.

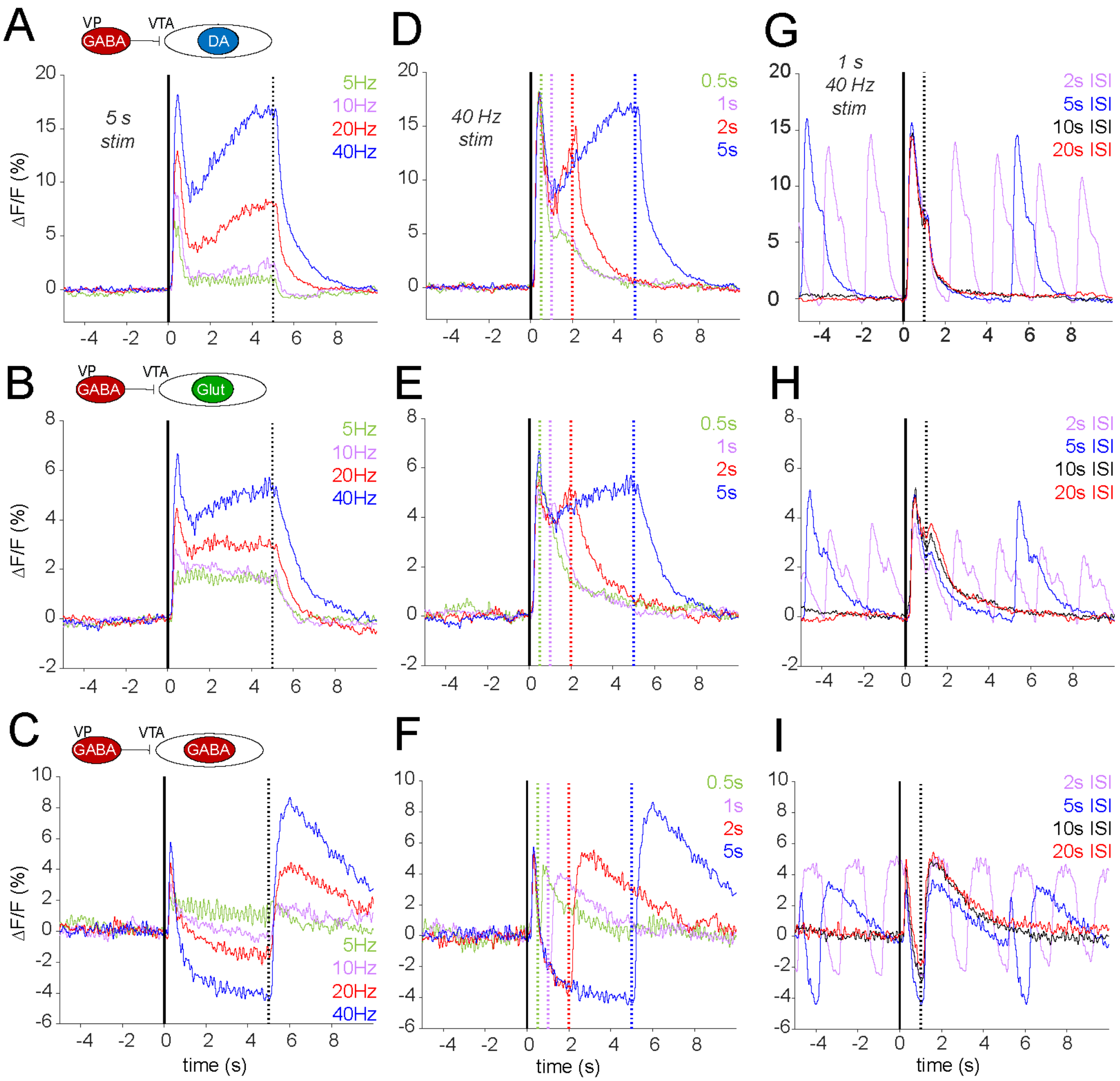

**Fig. S2. VTA cell-type responses to passive optogenetic stimulation of VP GABA neurons at different durations, frequencies and inter-stimulus intervals.** Responses (percent of ΔF/F) of GCaMP-expressing **A.** VTA DA, **B.** Glut, and **C.** GABA neurons to different frequencies of VP GABA neuron stim (t=0) delivered for 5 s with a 20s ISI. **D.** VTA DA, **E.** Glut, and **F.** GABA neuron responses to 40 Hz stim for different durations (onset at t=0s), with the end of each stim noted by a vertical line of the corresponding color and 20s ISI. **G.** VTA DA, **H.** Glut, and **I.** GABA neuron responses to 40-Hz 1-s stim with variable ISI. Note that with shorter ISIs signals have not returned to baseline prior to stim onset. Data are presented as mean. Related to Fig. 3.

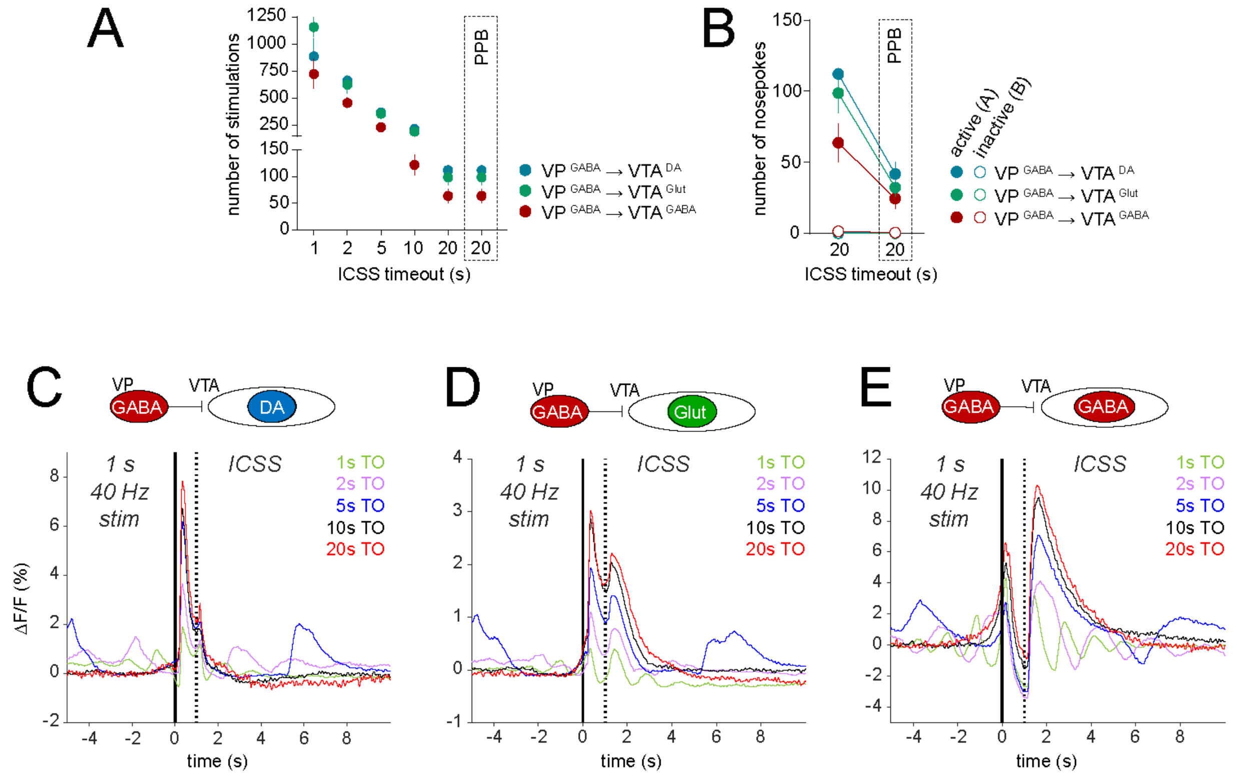

**Fig. S3. VTA cell-type responses during VP GABA neuron ICSS with different timeout periods. A.** Number of self-administered VP GABA neuron stims for each cohort of GCAMP recordings. An increasing timeout (TO) period was imposed between stim availability across daily sessions. The number of stims delivered during PPB is a replay of and thus identical to the 20s TO condition. **B.** Number of nosepokes made into the active and inactive holes during the 20-s TO ICSS session and subsequent PPB. **C.** VTA DA, **D.** Glut and **E.** GABA neuron GCaMP responses (percent of ΔF/F) to ICSS (t=0s) during the sessions with different TO periods imposed. Note that due to the high rate of ICSS, when TO periods were short GCaMP signals had not returned to baseline prior to stim onset. Data are presented as A & B, mean ± SEM; C-E, mean. Related to Fig. 4.

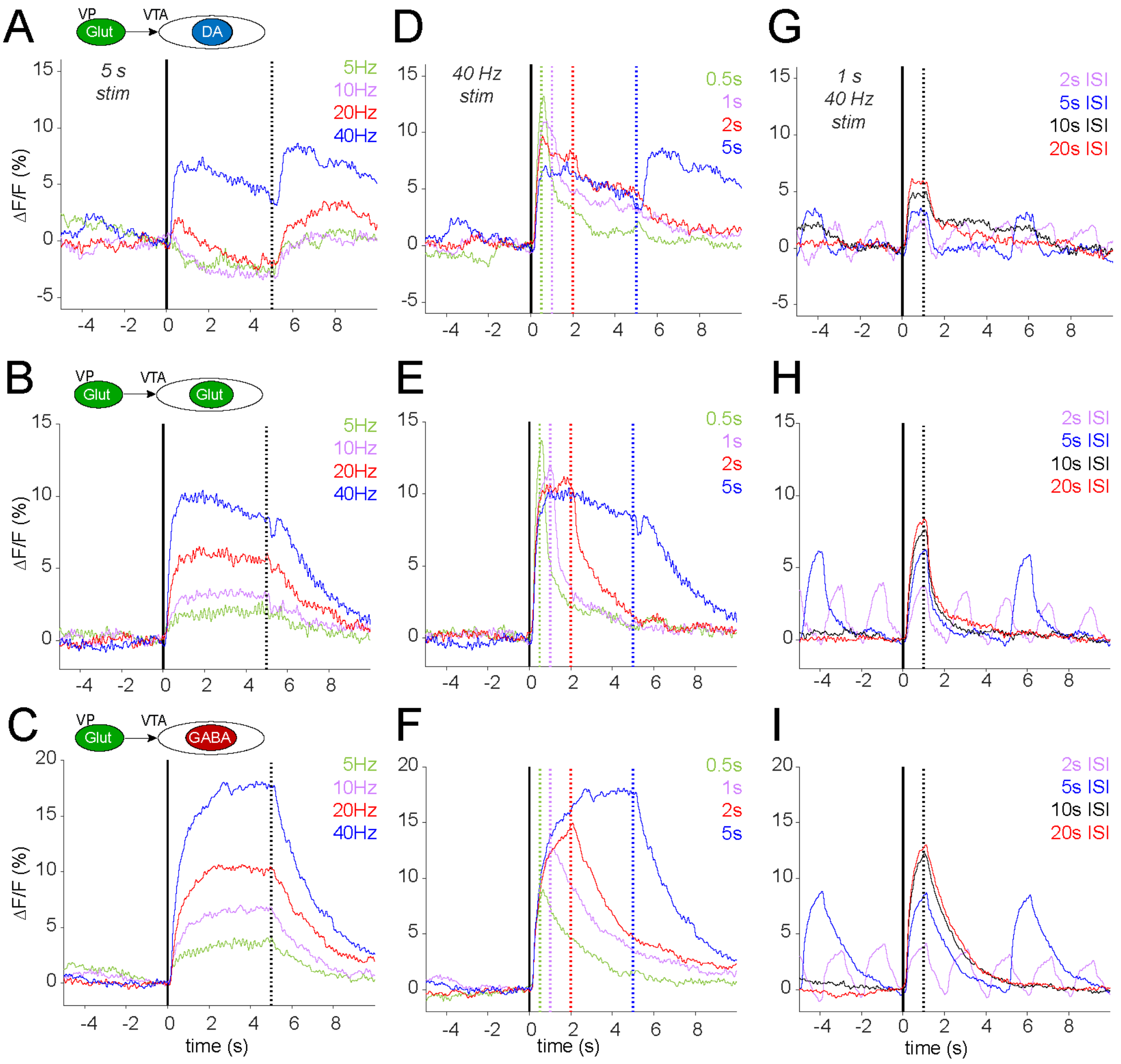

**Fig. S4. VTA cell-type responses to passive optogenetic stimulation of VP Glut neurons at different durations, frequencies and inter-stimulus intervals.** Responses (percent of ΔF/F) of GCaMP-expressing **A**. VTA DA, **B.** Glut, and **C.** GABA neurons to different frequencies of VP Glut neuron stim (t=0) delivered for 5 s with a 20s ISI. **D.** VTA DA, **E.** Glut, and **F.** GABA neuron responses to 40 Hz stim for different durations (onset at t=0s), with the end of each stim noted by a vertical line of the corresponding color and 20s ISI. **G.** VTA DA, **H.** Glut, and **I.** GABA neuron responses to 40-Hz 1-s stim with variable ISI. Note that with shorter ISIs signals have not returned to baseline prior to stim onset. Data are presented as mean. Related to Fig. 5.

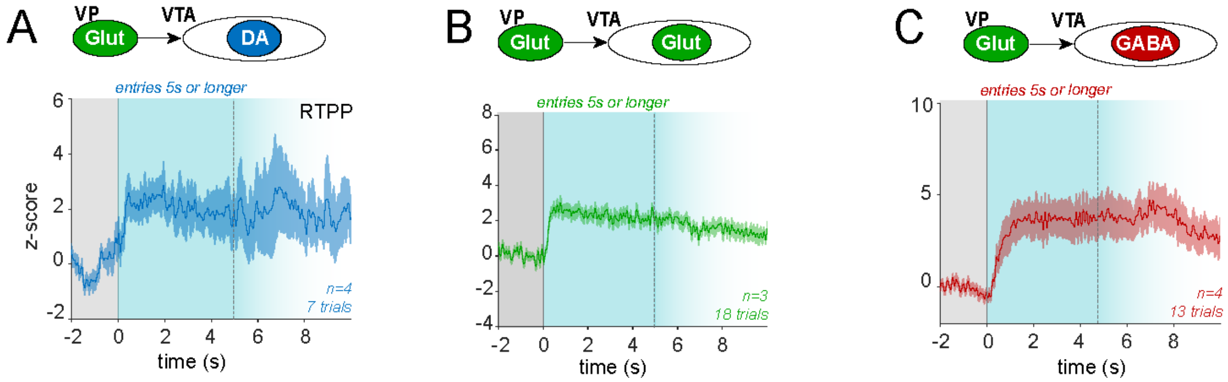

**Fig. S5. VTA cell-type responses to more sustained VP Glut neuron stimulation during RTPP assay.** In this experiment active side entry activates VP Glut neurons and because this is aversive mice make relatively few and relatively brief entries into the active side. However, some of the mice do make a few more sustained entries and these are the GCaMP responses (z-score) of **A.** VTA DA, **B.** VTA Glut, and **C.** VTA GABA neurons for entries sustained for $\geq$5s in the stim side (blue shading, fades after 5s) and preceded by $\geq$2s in the no-stim side (gray shading). Data are presented as mean ± SEM. Related to Fig. 5.

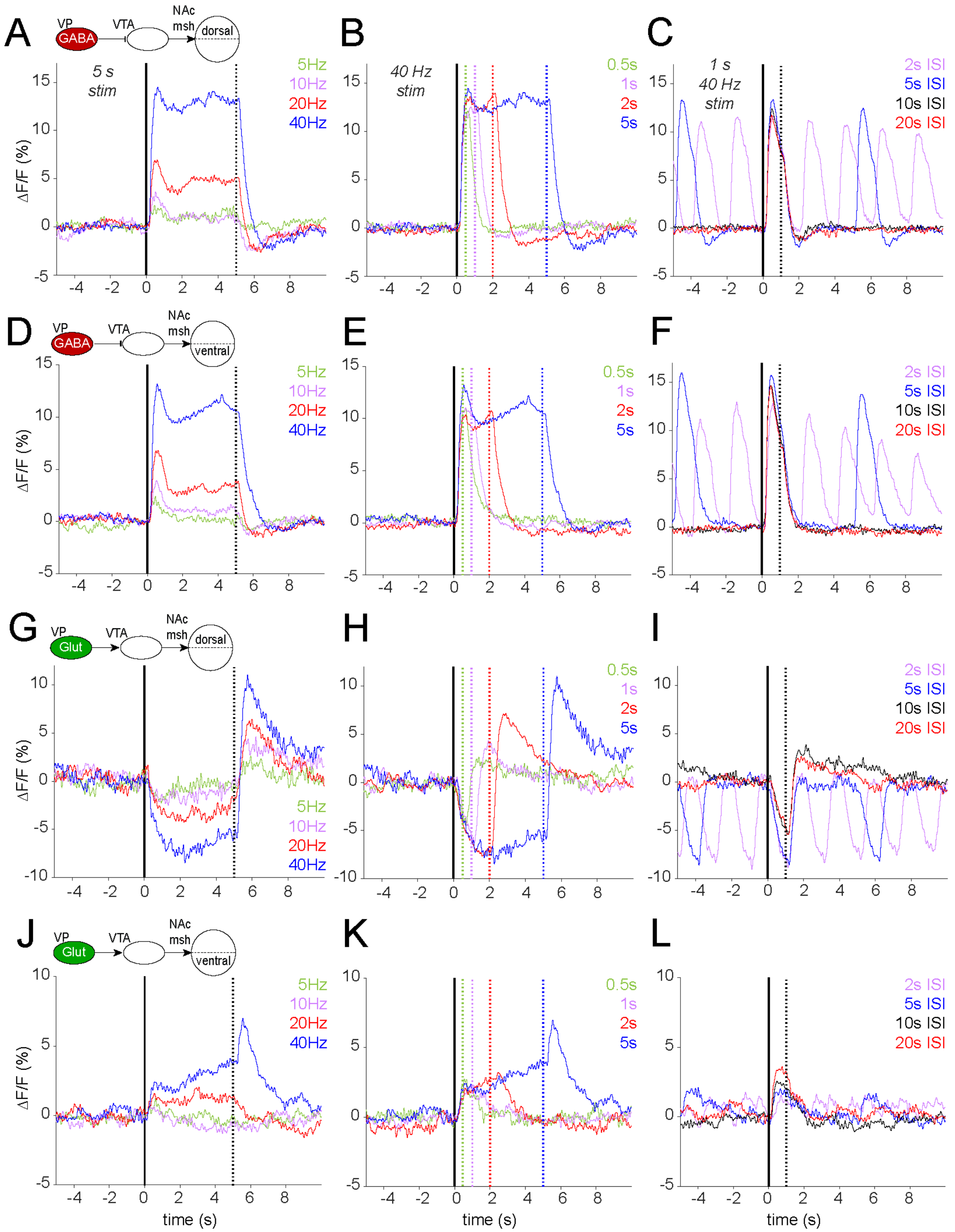

**Fig. S6. Dopamine release in the dorsal and ventral medial shell of NAc subregions in response to varied durations, frequencies and inter-stim intervals of optogenetic stimulation**

**of VP GABA and glutamate neuron terminals in VTA.** DA dLight responses (percent of ΔF/F) in dmsh of NAc to stimulation (onset t=0) of VP GABA terminals in VTA with varied **A.** frequency (fixed at 5s duration, 20s ISI), **B.** duration (fixed at 40Hz, 20s ISI), or **C.** ISI (fixed at 40Hz, 1s duration). DA dLight responses in vmsh of NAc to stimulation of VP GABA terminals in VTA with varied **D.** frequency (fixed at 5s duration, 20s ISI), **E.** duration (fixed at 40Hz, 20s ISI), or **F.** ISI (fixed at 40Hz, 1s duration). DA dLight responses in dmsh of NAc to stimulation of VP Glut terminals in VTA with varied **G.** frequency (fixed at 5s duration, 20s ISI), **H.** duration (fixed at 40Hz, 20s ISI), or **I.** ISI (fixed at 40Hz, 1s duration). DA dLight responses in dmsh NAc to stimulation (onset t=0) of VP GABA terminal in VTA with varied **J.** frequency (fixed at 5s duration, 20s ISI), **K.** duration (fixed at 40Hz, 20s ISI), or **L.** ISI (fixed at 40Hz, 1s duration). Data are presented as mean. Related to Fig. 6.

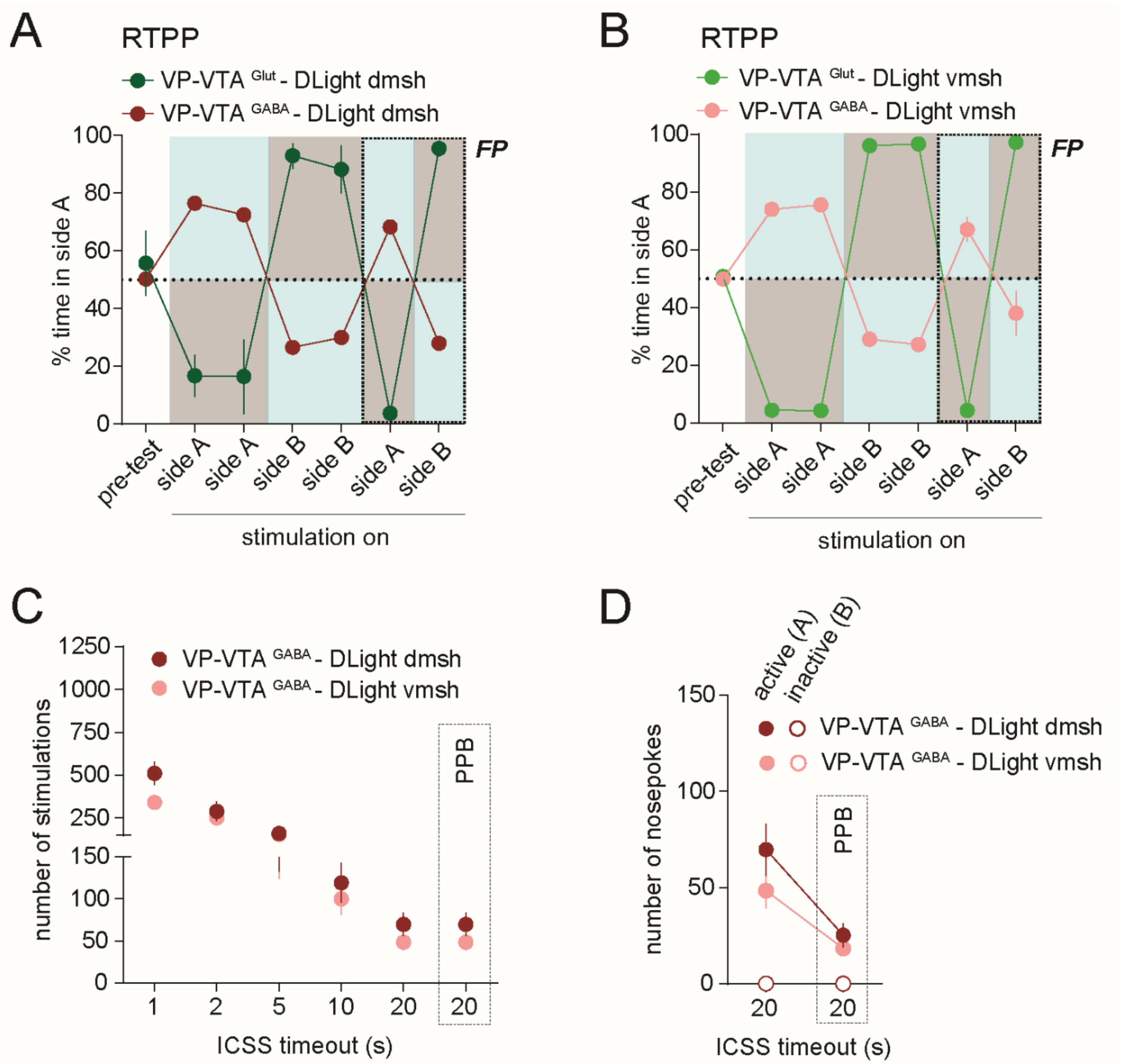

**Fig. S7. Behavioral responses to stimulation of VP GABA and Glut terminals in VTA.** Fraction of time spent in side A during RTPP for stim of VP Glut or GABA terminals in VTA, when recording DLight responses from **A**. NAc dmsh and **B.** NAc vmsh. Blue shading represents stimulated side; recordings were made only during the last two sessions. **C**. Number of nosepokes leading to self-stim of VP GABA terminals in VTA for each cohort of mice used in DLight recordings and for each timeout (TO) period tested. The number of stims delivered during passive playback (PPB) is a replay of and thus identical to the 20s TO condition. **D**. Number of nosepokes made into the active and inactive holes during the 20-s TO ICSS session and subsequent PPB session. Data are presented as mean ± SEM. Related to Fig. 6.

| Cell type / Pathway | Genotype | n= | Virus in VP | Virus in VTA | Virus in NAc |
|---|---|---|---|---|---|
| VP GABA | VGAT Cre/Cre | 6 | AAV5-DIO-GCaMP6f | | |
| VP Glut | VGLUT2 Cre/Cre | 7 | AAV5-DIO-GCaMP6f | | |
| VP GABA → VTA DA | VGAT +/FlpO ; DAT +/Cre | 5 | AAVDJ-fDIO-ChR2:YFP | AAV5-DIO-GCaMP6f | |
| VP GABA → VTA Glut | VGAT +/FlpO ; VGLUT2 +/Cre | 8 | AAVDJ-fDIO-ChR2:YFP | AAV5-DIO-GCaMP6f | |
| VP GABA → VTA GABA | VGAT Cre/Cre | 5 | AAV5-DIO-ChR2:mCherry | AAV5-DIO-GCaMP6f | |
| VP Glut → VTA DA | VGLUT2 +/Cre ; DAT +/FlpO | 6 | AAV5-DIO-ChR2:mCherry | AAVDJ-fDIO-GCaMP7f | |
| VP Glut → VTA Glut | VGLUT2 Cre/Cre | 7 | AAV5-DIO-ChR2:mCherry | AAV5-DIO-GCaMP6f | |
| VP Glut → VTA GABA | VGLUT2 +/Cre ; VGAT +/FlpO | 6 | AAV5-DIO-ChR2:mCherry | AAVDJ-fDIO-GCaMP7f | |
| VP-VTA GABA → NAc dmsh | VGAT Cre/Cre | 6 | AAV5-DIO-ChR2:mCherry | | AAV5-CAG-dLight1.1 |
| VP-VTA GABA → NAc vmsh | VGAT Cre/Cre | 5 | AAV5-DIO-ChR2:mCherry | | AAV5-CAG-dLight1.1 |
| VP-VTA Glut → NAc dmsh | VGLUT2 Cre/Cre | 4 | AAV5-DIO-ChR2:mCherry | | AAV5-CAG-dLight1.1 |
| VP-VTA Glut → NAc vmsh | VGLUT2 Cre/Cre | 4 | AAV5-DIO-ChR2:mCherry | | AAV5-CAG-dLight1.1 |

**Table S1. Combinatorial Cre- or FLP-dependent adeno-associated virus (AAV) vectors strategies to achieve expression of ChR2 in VP cell types, GCaMP in VTA cell types, and ubiquitous expression of dLight in NAc.** Related to Fig. 1 to 7.

| | | | | | variables | | | | values | | Statistical analysis | | | | | post-hoc test | | | |
|---|---|---|---|---|---|---|---|---|---|---|---|---|---|---|---|---|---|---|---|
| Figure | panel | experiment | VP cell type | VTA cell type | independent | dependent | n= | conditions / groups | mean | sem | type | factors | F or t value | p value | asterisk | type | comparison | p value | asterisk |
| 1 | | | | | | | | | | | | | | | | | | | |
| | A | HFHS | GABA | | first 10 | baseline | 6 | 3 | -0.01238 | 0.238 | RM 1-way ANOVA | time period | F(2,5) = 19.11 | 0.0049 | ** | Tukey | baseline vs. pre-event | 0.0098 | ** |
| | | CPP | | | | pre-event | | | 1.964 | 0.3012 | | | | | | | baseline vs. event | 0.0184 | * |
| | | | | | | event | | | 2.174 | 0.4685 | | | | | | | pre-event vs. event | 0.5259 | ns |
| | | | | | last 10 | baseline | 6 | 3 | -0.1387 | 0.2984 | RM 1-way ANOVA | time period | F(2,5)=36.74 | 0.0013 | ** | Tukey | baseline vs. pre-event | 0.0059 | ** |
| | | | | | | pre-event | | | 1.991 | 0.3017 | | | | | | | baseline vs. event | 0.0023 | ** |
| | | | | | | event | | | 2.113 | 0.3196 | | | | | | | pre-event vs. event | 0.4487 | ns |
| | | | | | event z-score | first 10 | 6 | 2 | 2.174 | 0.4685 | *t*-test | | t(5)= 0.1306 | 0.9012 | ns | | | | |
| | | | | | | last 10 | | | 2.113 | 0.3196 | | | | | | | | | |
| | B | HFHS | Glutamate | | first 10 | baseline | 7 | 3 | -0.4174 | 0.2068 | RM 1-way ANOVA | time period | F(2,6)=41.71 | < 0.0001 | **** | Tukey | baseline vs. pre-event | 0.001 | ** |
| | | CPP | | | | pre-event | | | 1.995 | 0.2569 | | | | | | | baseline vs. event | 0.0013 | ** |
| | | | | | | event | | | 1.856 | 0.2147 | | | | | | | pre-event vs. event | 0.7165 | ns |
| | | | | | last 10 | baseline | 7 | 3 | -0.4695 | 0.1588 | RM 1-way ANOVA | time period | F(2,6)=50.31 | < 0.0001 | **** | Tukey | baseline vs. pre-event | 0.0006 | *** |
| | | | | | | pre-event | | | 1.822 | 0.2092 | | | | | | | baseline vs. event | 0.0008 | *** |
| | | | | | | event | | | 1.731 | 0.1873 | | | | | | | pre-event vs. event | 0.838 | ns |
| | | | | | event z-score | first 10 | 7 | 2 | 1.856 | 0.2147 | *t*-test | | t(6)= 0.6419 | 0.5446 | ns | | | | |
| | | | | | | last 10 | | | 1.731 | 0.1873 | | | | | | | | | |
| | C | marble | GABA | | first 10 | baseline | 6 | 3 | 0.2136 | 0.2178 | RM 1-way ANOVA | time period | F(2,5)=27.55 | 0.0021 | ** | Tukey | baseline vs. pre-event | 0.0099 | ** |
| | | CPP | | | | pre-event | | | 2.192 | 0.3405 | | | | | | | baseline vs. event | 0.0053 | ** |
| | | | | | | event | | | 2.466 | 0.3485 | | | | | | | pre-event vs. event | 0.1357 | ns |
| | | | | | last 10 | baseline | 6 | 3 | -0.02067 | 0.2977 | RM 1-way ANOVA | time period | F(2,5)=11.18 | 0.0192 | * | Tukey | baseline vs. pre-event | 0.0455 | * |
| | | | | | | pre-event | | | 1.062 | 0.1473 | | | | | | | baseline vs. event | 0.0439 | * |
| | | | | | | event | | | 1.073 | 0.1511 | | | | | | | pre-event vs. event | 0.9705 | ns |
| | | | | | event z-score | first 10 | 6 | 2 | 2.466 | 0.3485 | *t*-test | | t(5)= 3.087 | 0.0273 | * | | | | |
| | | | | | | last 10 | | | 1.073 | 0.1511 | | | | | | | | | |
| | D | marble | Glutamate | | first 10 | baseline | 6 | 3 | -0.145 | 0.2795 | RM 1-way ANOVA | time period | F(2,5)=12.02 | 0.014 | * | Tukey | baseline vs. pre-event | 0.0419 | * |
| | | CPP | | | | pre-event | | | 1.506 | 0.3078 | | | | | | | baseline vs. event | 0.0314 | * |
| | | | | | | event | | | 1.427 | 0.2784 | | | | | | | pre-event vs. event | 0.8413 | ns |
| | | | | | last 10 | baseline | 6 | 3 | -0.6206 | 0.07159 | RM 1-way ANOVA | time period | F(2,5)= 7.457 | 0.0253 | * | Tukey | baseline vs. pre-event | 0.0135 | * |
| | | | | | | pre-event | | | 0.7118 | 0.2568 | | | | | | | baseline vs. event | 0.1736 | ns |
| | | | | | | event | | | 0.3867 | 0.4229 | | | | | | | pre-event vs. event | 0.5459 | ns |
| | | | | | event z-score | first 10 | 6 | 2 | 1.427 | 0.2784 | *t*-test | | t(5)=5.003 | 0.0041 | ** | | | | |
| | | | | | | last 10 | | | 0.3867 | 0.4229 | | | | | | | | | |
| | E | milk licking | GABA | | 20s ILI | baseline | 6 | 3 | 0.2466 | 0.2444 | RM 1-way ANOVA | time period | F(2,5)=11.47 | 0.0048 | ** | Tukey | baseline vs. pre-event | 0.0406 | * |
| | | | | | | pre-event | | | 1.884 | 0.3078 | | | | | | | baseline vs. event | 0.0251 | * |
| | | | | | | event | | | 2.588 | 0.4967 | | | | | | | pre-event vs. event | 0.2979 | ns |
| | | | | | event z-score | 1s ILI | 6 | 5 | 0.9839 | 0.1092 | RM 1-way ANOVA | ILI | F(4,5)= 7.575 | 0.0373 | * | | | | |
| | | | | | | 2s ILI | | | 1.388 | 0.08333 | | | | | | | | | |
| | | | | | | 5s ILI | | | 2.07 | 0.177 | | | | | | | | | |
| | | | | | | 10s ILI | | | 2.479 | 0.3642 | | | | | | | | | |
| | | | | | | 20s ILI | | | 2.588 | 0.4967 | | | | | | | | | |
| | | | | | event z-score | 1, 2, 5, 10, 20s ILI | 6 | 5 | | | linear regression | $R^2 = 0.3431$ | F(1,28)= 14.62 | 0.0007 | *** | | | | |
| | | | | | event z-score | 1, 2, 5, 10, 20s ILI | animal 1 | 5 | | | linear regression | $R^2 = 0.8262$ | F(1,3)= 14.26 | 0.0325 | * | | | | |
| | | | | | event z-score | 1, 2, 5, 10, 20s ILI | animal 2 | 5 | | | linear regression | $R^2 = 0.8216$ | F(1,3)= 13.81 | 0.0339 | * | | | | |
| | | | | | event z-score | 1, 2, 5, 10, 20s ILI | animal 3 | 5 | | | linear regression | $R^2 = 0.8271$ | F(1,3)= 14.35 | 0.0323 | * | | | | |
| | | | | | event z-score | 1, 2, 5, 10, 20s ILI | animal 4 | 5 | | | linear regression | $R^2 = 0.5798$ | F(1,3)= 4.139 | 0.1348 | ns | | | | |
| | | | | | event z-score | 1, 2, 5, 10, 20s ILI | animal 5 | 5 | | | linear regression | $R^2 = 0.2297$ | F(1,3)= 0.8946 | 0.414 | ns | | | | |
| | | | | | event z-score | 1, 2, 5, 10, 20s ILI | animal 6 | 5 | | | linear regression | $R^2 = 0.7964$ | F(1,3)= 11.73 | 0.0417 | * | | | | |
| | F | milk licking | Glutamate | | 20s ILI | baseline | 6 | 3 | 0.2862 | 0.3266 | RM 1-way ANOVA | time period | F(2,5)=17.13 | 0.0042 | ** | Tukey | baseline vs. pre-event | 0.0056 | ** |
| | | | | | | pre-event | | | 2.283 | 0.2498 | | | | | | | baseline vs. event | 0.0193 | * |
| | | | | | | event | | | 2.837 | 0.4414 | | | | | | | pre-event vs. event | 0.3676 | ns |
| | | | | | event z-score | 1s ILI | 6 | 5 | 0.9987 | 0.1403 | RM 1-way ANOVA | ILI | F(4,5)=13.71 | 0.005 | ** | | | | |
| | | | | | | 2s ILI | | | 1.42 | 0.1675 | | | | | | | | | |
| | | | | | | 5s ILI | | | 1.97 | 0.3186 | | | | | | | | | |
| | | | | | | 10s ILI | | | 2.428 | 0.429 | | | | | | | | | |
| | | | | | | 20s ILI | | | 2.837 | 0.4414 | | | | | | | | | |
| | | | | | event z-score | 1, 2, 5, 10, 20s ILI | 6 | 5 | | | linear regression | $R^2 = 0.39$ | F(1,28)= 17.90 | 0.0002 | *** | | | | |
| | | | | | event z-score | 1, 2, 5, 10, 20s ILI | animal 1 | 5 | | | linear regression | $R^2 = 0.9235$ | F(1,3)= 36.19 | 0.0092 | ** | | | | |
| | | | | | event z-score | 1, 2, 5, 10, 20s ILI | animal 2 | 5 | | | linear regression | $R^2 = 0.5383$ | F(1,3)= 3.498 | 0.1582 | ns | | | | |
| | | | | | event z-score | 1, 2, 5, 10, 20s ILI | animal 3 | 5 | | | linear regression | $R^2 = 0.8650$ | F(1,3)= 19.22 | 0.022 | * | | | | |
| | | | | | event z-score | 1, 2, 5, 10, 20s ILI | animal 4 | 5 | | | linear regression | $R^2 = 0.9503$ | F(1,3)= 57.40 | 0.0048 | ** | | | | |
| | | | | | event z-score | 1, 2, 5, 10, 20s ILI | animal 5 | 5 | | | linear regression | $R^2 = 0.6366$ | F(1,3)= 5.256 | 0.1057 | ns | | | | |
| | | | | | event z-score | 1, 2, 5, 10, 20s ILI | animal 6 | 5 | | | linear regression | $R^2 = 0.9638$ | F(1,3)= 79.82 | 0.003 | ** | | | | |
| 2 | | | | | | | | | | | | | | | | | | | |
| | A | choco vs. quinine | GABA | | chocolate | baseline | 6 | 3 | 0.0104 | 0.3785 | RM 1-way ANOVA | time period | F(2,5)=13.42 | 0.0143 | * | Tukey | baseline vs. pre-event | 0.0492 | * |
| | | pellets | | | | pre-event | | | 1.841 | 0.3064 | | | | | | | baseline vs. event | 0.0239 | * |
| | | | | | | event | | | 2.22 | 0.3065 | | | | | | | pre-event vs. event | 0.0007 | *** |
| | | | | | quinine | baseline | 6 | 3 | -0.2326 | 0.3249 | RM 1-way ANOVA | time period | F(2,5)=4.435 | 0.0611 | ns | | | | |
| | | | | | | pre-event | | | 0.3845 | 0.495 | | | | | | | | | |
| | | | | | | event | | | 1.309 | 0.1381 | | | | | | | | | |
| | | | | | event z-score | chocolate | 6 | 2 | 2.22 | 0.3065 | *t*-test | | t(5)= 4.554 | 0.0061 | ** | | | | |
| | | | | | | qunine | | | 1.309 | 0.1381 | | | | | | | | | |
| | B | choco vs. quinine | Glutamate | | chocolate | baseline | 6 | 3 | -0.2016 | 0.2592 | RM 1-way ANOVA | time period | F(2,5)=10.94 | 0.0211 | * | Tukey | baseline vs. pre-event | 0.0484 | * |
| | | pellets | | | | pre-event | | | 2.215 | 0.5372 | | | | | | | baseline vs. event | 0.046 | * |
| | | | | | | event | | | 2.287 | 0.555 | | | | | | | pre-event vs. event | 0.2364 | ns |
| | | | | | quinine | baseline | 6 | 3 | -0.6277 | 0.2014 | RM 1-way ANOVA | time period | F(2,5) = 20.77 | 0.0052 | ** | Tukey | baseline vs. pre-event | 0.008 | ** |
| | | | | | | pre-event | | | 1.785 | 0.3546 | | | | | | | baseline vs. event | 0.0192 | * |
| | | | | | | event | | | 1.73 | 0.4719 | | | | | | | pre-event vs. event | 0.9129 | ns |
| | | | | | event z-score | chocolate | 6 | 2 | 2.287 | 0.555 | *t*-test | | t(5) = 0.8759 | 0.4212 | ns | | | | |
| | | | | | | qunine | | | 1.73 | 0.4719 | | | | | | | | | |
| | C | Shock | GABA | | stimulus | cue - conditioning | 5 | 4 | 3.124 | 0.6821 | RM 2-way ANOVA | stimulus type | F (1, 4) = 4.501 | 0.1012 | ns | Sidak | conditioning vs. extinction - cue | 0.7592 | ns |
| | | | | | | cue - extinction | | | 4.362 | 0.6501 | | session | F (1, 4) = 12.23 | 0.025 | * | | conditioning vs. extinction - shock | 0.0069 | ** |
| | | | | | | shock - conditioning | | | 12.36 | 2.569 | | interaction | F (1, 4) = 23.95 | 0.0081 | ** | | | | |
| | | | | | | shock - extinction | | | 1.759 | 0.7314 | | | | | | | | | |
| | D | Shock | Glutamate | | stimulus | cue - conditioning | 4 | 4 | 0.09528 | 0.4692 | RM 2-way ANOVA | stimulus type | F (1, 3) = 6.542 | 0.0834 | ns | Sidak | conditioning vs. extinction - cue | 0.6343 | ns |
| | | | | | | cue - extinction | | | -0.509 | 0.4068 | | session | F (1, 3) = 68.18 | 0.0037 | ** | | conditioning vs. extinction - shock | 0.0173 | * |
| | | | | | | shock - conditioning | | | 3.32 | 0.6499 | | interaction | F (1, 3) = 13.25 | 0.0357 | * | | | | |
| | | | | | | shock - extinction | | | -0.4275 | 0.3298 | | | | | | | | | |
| 3 | | | | | | | | | | | | | | | | | | | |
| | B | RTPP | GABA | DA | stim | test day | 5 | GABA --> DA | | | RM 2-way ANOVA | group | F (2, 15) = 0.2964 | 0.7478 | ns | Sidak | GABA --> DA | | |
| | | | | Glut | | | | pre-test | 48.206 | 4.086964 | | days | F (6, 90) = 389.4 | < 0.0001 | **** | | pre-test vs. test 1 | < 0.0001 | **** |
| | | | | GABA | | | | test 1 | 90.224 | 4.7614 | | interaction | F (12, 90) = 2.690 | 0.0039 | ** | | pre-test vs. test 2 | < 0.0001 | **** |
| | | | | | | | | test 2 | 88.432 | 3.907386 | | | | | | | pre-test vs. test 3 | < 0.0001 | **** |
| | | | | | | | | test 3 | 13.692 | 1.86437 | | | | | | | pre-test vs. test 4 | < 0.0001 | **** |
| | | | | | | | | test 4 | 10.652 | 2.774894 | | | | | | | pre-test vs. test 5 | < 0.0001 | **** |
| | | | | | | | | test 5 | 83.276 | 0.7925006 | | | | | | | pre-test vs. test 6 | < 0.0001 | **** |
| | | | | | | | | test 6 | 22.552 | 5.726597 | | | | | | | | | |
| | | | | | | | 8 | GABA --> Glut | | | | | | | | | GABA --> Glut | | |
| | | | | | | | | pre-test | 50.6125 | 3.507871 | | | | | | | pre-test vs. test 1 | < 0.0001 | **** |
| | | | | | | | | test 1 | 93.025 | 1.675426 | | | | | | | pre-test vs. test 2 | < 0.0001 | **** |
| | | | | | | | | test 2 | 94.1125 | 1.282637 | | | | | | | pre-test vs. test 3 | < 0.0001 | **** |
| | | | | | | | | test 3 | 8.1 | 1.848841 | | | | | | | pre-test vs. test 4 | < 0.0001 | **** |
| | | | | | | | | test 4 | 6.925 | 1.508162 | | | | | | | pre-test vs. test 5 | < 0.0001 | **** |
| | | | | | | | | test 5 | 90.5375 | 1.777532 | | | | | | | pre-test vs. test 6 | < 0.0001 | **** |

| | | | | | | | | | | | | | | | | | | | |
|---|---|---|---|---|---|---|---|---|---|---|---|---|---|---|---|---|---|---|---|
| | | | | | | | | test6 | 9.1 | 1.764835 | | | | | | | | | |
| | | | | | | | 5 | GABA --> GABA | | | | | | | | | GABA --> GABA | | |
| | | | | | | | | pre-test | 48.8 | 7.186098 | | | | | | | pre-test vs. test1 | < 0.0001 | **** |
| | | | | | | | | test1 | 85.6 | 4.261456 | | | | | | | pre-test vs. test2 | < 0.0001 | **** |
| | | | | | | | | test2 | 85.8 | 4.651882 | | | | | | | pre-test vs. test3 | < 0.0001 | **** |
| | | | | | | | | test3 | 14 | 4.949748 | | | | | | | pre-test vs. test4 | < 0.0001 | **** |
| | | | | | | | | test4 | 14.2 | 4.841487 | | | | | | | pre-test vs. test5 | < 0.0001 | **** |
| | | | | | | | | test5 | 76.6 | 2.942788 | | | | | | | pre-test vs. test6 | < 0.0001 | **** |
| | | | | | | | | test6 | 20 | 3.754997 | | | | | | | | | |
| | D | Passive stim | GABA | DA | stim | pre | 5 | 3 | -0.1112 | 0.3174 | RM 1-way ANOVA | time period | F(2,4)=41.37 | 0.0011 | ** | Tukey | pre vs. early | 0.0071 | ** |
| | | | | | | early | | | 18.13 | 2.685 | | | | | | | pre vs. late | 0.0039 | ** |
| | | | | | | late | | | 17.48 | 2.23 | | | | | | | early vs. late | 0.8668 | ns |
| | E | Passive stim | GABA | Glut | stim | pre | 8 | 3 | 0.4823 | 0.1606 | RM 1-way ANOVA | time period | F(2,7)=27.81 | 0.0001 | *** | Tukey | pre vs. early | 0.0003 | *** |
| | | | | | | early | | | 7.873 | 1.006 | | | | | | | pre vs. late | 0.0052 | ** |
| | | | | | | late | | | 6.841 | 1.374 | | | | | | | early vs. late | 0.5042 | ns |
| | F | Passive stim | GABA | GABA | stim | pre | 5 | 3 | 0.1802 | 0.3742 | RM 1-way ANOVA | time period | F(2,4)=9.374 | 0.037 | * | Tukey | pre vs. early | 0.2788 | ns |
| | | | | | | early | | | 3.173 | 1.763 | | | | | | | pre vs. late | 0.0001 | *** |
| | | | | | | late | | | -2.821 | 0.5015 | | | | | | | early vs. late | 0.0541 | ns |
| | H | RTPP 5s | GABA | DA | stim | pre | 5 | 3 | -0.614 | 0.2463 | RM 1-way ANOVA | time period | F(2,4)=31.45 | 0.0033 | ** | Tukey | pre vs. early | 0.0069 | ** |
| | | | | | | early | | | 7.925 | 1.225 | | | | | | | pre vs. late | 0.0012 | ** |
| | | | | | | late | | | 3.182 | 0.3406 | | | | | | | early vs. late | 0.0402 | * |
| | I | RTPP 5s | GABA | Glut | stim | pre | 8 | 3 | 0.3145 | 0.2774 | RM 1-way ANOVA | time period | F(2,7)=17.37 | 0.0003 | *** | Tukey | pre vs. early | 0.0015 | ** |
| | | | | | | early | | | 3.666 | 0.5539 | | | | | | | pre vs. late | 0.9982 | ns |
| | | | | | | late | | | 0.3577 | 0.7292 | | | | | | | early vs. late | 0.0025 | ** |
| | J | RTPP 5s | GABA | GABA | stim | pre | 5 | 3 | 0.5119 | 0.07927 | RM 1-way ANOVA | time period | F(2,4)=64.96 | < 0.0001 | **** | Tukey | pre vs. early | 0.003 | ** |
| | | | | | | early | | | -1.307 | 0.2476 | | | | | | | pre vs. late | 0.0015 | ** |
| | | | | | | late | | | -2.037 | 0.2638 | | | | | | | early vs. late | 0.038 | * |
| | K | Passive vs. | GABA | DA | stim | passive - early | 5 | 4 | 18.13071 | 2.684644 | RM 2-way ANOVA | time period | F (1, 4) = 8.370 | p = 0.0444 | * | Sidak | passive vs. RTPP - early | 0.0019 | ** |
| | | RTPP | | | | RTPP - early | | | 7.925275 | 1.22531 | | test | F (1, 4) = 30.51 | p = 0.0052 | ** | | passive vs. RTPP - late | 0.0005 | *** |
| | | | | | | passive - late | | | 17.48433 | 2.2299 | | interaction | F (1, 4) = 6.169 | p = 0.0679 | ns | | | | |
| | | | | | | RTPP - late | | | 3.182342 | 0.3406029 | | | | | | | | | |
| | L | Passive vs. | GABA | Glut | stim | passive - early | 8 | 3 | 7.873426 | 1.005671 | RM 2-way ANOVA | time period | F (1, 7) = 12.59 | p = 0.0094 | ** | Sidak | passive vs. RTPP - early | 0.0006 | *** |
| | | RTPP | | | | RTPP - early | | | 3.665979 | 0.5538577 | | assay type | F (1, 7) = 38.28 | p = 0.0005 | *** | | passive vs. RTPP - late | < 0.0001 | **** |
| | | | | | | passive - late | | | 6.840797 | 1.374191 | | interaction | F (1, 7) = 6.456 | p = 0.0386 | * | | | | |
| | | | | | | RTPP - late | | | 0.3576861 | 0.7291539 | | | | | | | | | |
| | M | Passive vs. | GABA | GABA | stim | passive - early | 5 | 3 | 3.173058 | 1.762514 | RM 2-way ANOVA | time period | F (1, 4) = 13.71 | p = 0.0208 | * | Sidak | passive vs. RTPP - early | 0.0367 | * |
| | | RTPP | | | | RTPP - early | | | -1.306934 | 0.2476216 | | assay type | F (1, 4) = 4.772 | p = 0.0943 | ns | | passive vs. RTPP - late | 0.7873 | ns |
| | | | | | | passive - late | | | -2.820807 | 0.5015188 | | interaction | F (1, 4) = 10.16 | p = 0.0333 | * | | | | |
| | | | | | | RTPP - late | | | -2.036961 | 0.2638479 | | | | | | | | | |
| 4 | | | | | | | | | | | | | | | | | | | |
| | B | Passive | GABA | DA | stim | Passive | 5 | 3 | 14.62 | 2.363 | RM 1-way ANOVA | assay type | F(2,4)= 28.93 | 0.0009 | *** | Tukey | passive stim vs. ICSS | 0.0071 | ** |
| | | ICSS | | | | ICSS | | | 6.081 | 1.611 | | | | | | | passive stim vs. PPB | 0.0324 | * |
| | | PPB | | | | PPB | | | 10.06 | 2.097 | | | | | | | ICSS vs. PPB | 0.0195 | * |
| | C | Passive | GABA | DA | pre-event | Passive | 5 | 4 | -0.08585 | 0.233 | RM 1-way ANOVA | assay type | F(3,4)= 2.312 | 0.1927 | ns | | | | |
| | | ICSS | | | | ICSS | | | 0.2172 | 0.1145 | | | | | | | | | |
| | | PPB | | | | PPB | | | -0.2182 | 0.01446 | | | | | | | | | |
| | | nosepoke PPB | | | | nosepoke PPB | | | 0.3043 | 0.1385 | | | | | | | | | |
| | D | Passive | GABA | Glut | stim | Passive | 8 | 3 | 6.172 | 0.7279 | RM 1-way ANOVA | assay type | F(2,7)= 9.567 | 0.0087 | ** | Tukey | passive stim vs. ICSS | 0.0694 | ns |
| | | ICSS | | | | ICSS | | | 4.137 | 0.4967 | | | | | | | passive stim vs. PPB | 0.8394 | ns |
| | | PPB | | | | PPB | | | 6.457 | 0.6545 | | | | | | | ICSS vs. PPB | 0.0027 | ** |
| | E | Passive | GABA | Glut | pre-event | Passive | 8 | 4 | -0.04639 | 0.2736 | RM 1-way ANOVA | assay type | F(3,7)= 3.894 | 0.0622 | ns | | | | |
| | | ICSS | | | | ICSS | | | 0.3883 | 0.2085 | | | | | | | | | |
| | | PPB | | | | PPB | | | -0.02923 | 0.1201 | | | | | | | | | |
| | | nosepoke PPB | | | | nosepoke PPB | | | 0.7483 | 0.1214 | | | | | | | | | |
| | F | Passive | GABA | GABA | stim | Passive - max | 5 | 6 | 3.089698 | 0.8590032 | RM 2-way ANOVA | assay type | F(2,8)= 0.004358 | 0.9957 | ns | Sidak | passive vs. ICSS - max | 0.8846 | ns |
| | | ICSS | | | | ICSS - max | | | 2.597996 | 0.4808929 | | peak type | F(1,4)= 101.5 | 0.0005 | *** | | passive vs. PPB - max | 0.8063 | ns |
| | | PPB | | | | PPB max | | | 2.4808 | 0.7295148 | | interaction | F(2,8)= 0.7544 | 0.501 | ns | | ICSS vs. PPB - max | 0.998 | ns |
| | | | | | | Passive - min | | | -1.266482 | 0.3155479 | | | | | | | passive vs. ICSS - min | 0.871 | ns |
| | | | | | | ICSS - min | | | -0.7525226 | 0.2920639 | | | | | | | passive vs. PPB - min | 0.8582 | ns |
| | | | | | | PPB min | | | -0.7324064 | 0.3025935 | | | | | | | ICSS vs. PPB - min | > 0.9999 | ns |
| | G | Passive | GABA | GABA | pre-event | Passive | 5 | 4 | -0.2304 | 0.3738 | RM 1-way ANOVA | assay type | F(3,4)= 16.59 | 0.0016 | ** | Tukey | passive stim vs. ICSS | 0.0282 | * |
| | | ICSS | | | | ICSS | | | 1.723 | 0.2802 | | | | | | | passive stim vs. PPB | > 0.9999 | ns |
| | | PPB | | | | PPB | | | -0.213 | 0.1065 | | | | | | | passive stim vs. nosepoke PPB | 0.0656 | ns |
| | | nosepoke PPB | | | | nosepoke PPB | | | 1.543 | 0.3468 | | | | | | | ICSS vs. PPB | 0.0166 | * |
| | | | | | | | | | | | | | | | | | ICSS vs. nosepoke PPB | 0.8442 | ns |
| | | | | | | | | | | | | | | | | | PPB vs. nosepoke PPB | 0.0451 | * |
| 5 | | | | | | | | | | | | | | | | | | | |
| | B | RTPP | Glut | DA | stim | test day | 6 | Glut --> DA | | | RM 2-way ANOVA | group | F (2, 16) = 1.094 | 0.3587 | ns | Sidak | Glut --> DA | | |
| | | | | Glut | | | | pre-test | 51.87667 | 7.620903 | | days | F (6, 96) = 265.6 | < 0.0001 | **** | | pre-test vs. test1 | < 0.0001 | **** |
| | | | | GABA | | | | test1 | 4.218333 | 1.211746 | | interaction | F (12, 96) = 0.4224 | 0.9512 | ns | | pre-test vs. test2 | < 0.0001 | **** |
| | | | | | | | | test2 | 1.59 | 0.6387174 | | | | | | | pre-test vs. test3 | 0.0005 | *** |
| | | | | | | | | test3 | 79.73666 | 12.66596 | | | | | | | pre-test vs. test4 | < 0.0001 | **** |
| | | | | | | | | test4 | 94.295 | 2.566612 | | | | | | | pre-test vs. test5 | < 0.0001 | **** |
| | | | | | | | | test5 | 2.601667 | 0.6160975 | | | | | | | pre-test vs. test6 | < 0.0001 | **** |
| | | | | | | | | test6 | 96.41334 | 1.691131 | | | | | | | | | |
| | | | | | | | 7 | Glut --> Glut | | | | | | | | | Glut --> Glut | | |
| | | | | | | | | pre-test | 50.92857 | 7.561984 | | | | | | | pre-test vs. test1 | < 0.0001 | **** |
| | | | | | | | | test1 | 9.552857 | 4.16445 | | | | | | | pre-test vs. test2 | < 0.0001 | **** |
| | | | | | | | | test2 | 3.612857 | 2.001954 | | | | | | | pre-test vs. test3 | < 0.0001 | **** |
| | | | | | | | | test3 | 94.66428 | 2.174791 | | | | | | | pre-test vs. test4 | < 0.0001 | **** |
| | | | | | | | | test4 | 95.10286 | 2.26694 | | | | | | | pre-test vs. test5 | < 0.0001 | **** |
| | | | | | | | | test5 | 6.434286 | 3.4058 | | | | | | | pre-test vs. test6 | < 0.0001 | **** |
| | | | | | | | | test6 | 96.04857 | 1.980424 | | | | | | | | | |
| | | | | | | | 6 | Glut --> GABA | | | | | | | | | Glut --> GABA | | |
| | | | | | | | | pre-test | 53.33333 | 7.722981 | | | | | | | pre-test vs. test1 | < 0.0001 | **** |
| | | | | | | | | test1 | 8.166667 | 1.740051 | | | | | | | pre-test vs. test2 | < 0.0001 | **** |
| | | | | | | | | test2 | 7.833333 | 2.535306 | | | | | | | pre-test vs. test3 | < 0.0001 | **** |
| | | | | | | | | test3 | 87.16666 | 7.726218 | | | | | | | pre-test vs. test4 | < 0.0001 | **** |
| | | | | | | | | test4 | 94.83334 | 2.300966 | | | | | | | pre-test vs. test5 | < 0.0001 | **** |
| | | | | | | | | test5 | 4.166667 | 1.194897 | | | | | | | pre-test vs. test6 | < 0.0001 | **** |
| | | | | | | | | test6 | 95.16666 | 1.887091 | | | | | | | | | |
| | D | Passive 1s | Glut | DA | stim | pre | 6 | 2 | -0.3847 | 0.2389 | *t-test* | | t(5)= 13.75 | <0.0001 | **** | | | | |
| | | | | | | stim | | | 4.251 | 0.2664 | | | | | | | | | |
| | | Passive 5s | Glut | DA | stim | pre | 6 | 3 | -0.5346 | 0.07234 | RM 1-way ANOVA | time period | F(2,5)= 42.70 | 0.0001 | *** | Tukey | pre vs. early | 0.0001 | *** |
| | | | | | | early | | | 2.26 | 0.1851 | | | | | | | pre vs. late | 0.0027 | ** |
| | | | | | | late | | | 1.73 | 0.3049 | | | | | | | early vs. late | 0.4172 | ns |
| | E | Passive 1s | Glut | Glut | stim | pre | 7 | 2 | -0.2857 | 0.2894 | *t-test* | | t(6)= 7.787 | 0.0002 | *** | | | | |
| | | | | | | stim | | | 6.666 | 0.9175 | | | | | | | | | |
| | | Passive 5s | Glut | Glut | stim | pre | 7 | 3 | 0.3476 | 0.2792 | RM 1-way ANOVA | time period | F(2,6)= 50.48 | < 0.0001 | **** | Tukey | pre vs. early | 0.0006 | *** |
| | | | | | | early | | | 5.638 | 0.555 | | | | | | | pre vs. late | 0.0006 | *** |
| | | | | | | late | | | 5.234 | 0.4828 | | | | | | | early vs. late | 0.566 | ns |
| | F | Passive 1s | Glut | GABA | stim | pre | 6 | 2 | -0.3749 | 0.2552 | *t-test* | | t(5)= 5.921 | 0.002 | ** | | | | |

| | | | | | | | | | | | | | | | | | | | |
|---|---|---|---|---|---|---|---|---|---|---|---|---|---|---|---|---|---|---|---|
| | | | | | | stim | | | 6.882 | 1.276 | | | | | | | | | |
| | | Passive 5s | Glut | GABA | stim | pre<br>early<br>late | 6 | 3 | -0.1586<br>7.773<br>10.36 | 0.3228<br>0.6998<br>1.373 | RM 1-way ANOVA | time period | F(2,5)= 73.89 | 0.0002 | *** | Tukey | pre vs. early<br>pre vs. late<br>early vs. late | < 0.0001<br>0.0009<br>0.0507 | ****<br>***<br>ns |
| | G | RTPP 1s on | Glut | DA | stim onset | pre<br>stim | 6 | 3 | 0.3835<br>2.524 | 0.4931<br>0.2273 | *t*-test | | t(5)= 3.841 | 0.0121 | * | | | | |
| | H | RTPP 1s on | Glut | Glut | stim onset | pre<br>stim | 7 | 3 | 0.6625<br>5.414 | 0.5695<br>0.8113 | *t*-test | | t(6)= 8.411 | 0.0002 | *** | | | | |
| | I | RTPP 1s on | Glut | GABA | stim onset | pre<br>stim | 6 | 3 | 0.02325<br>4.909 | 0.3984<br>1.228 | *t*-test | | t(5)= 4.652 | 0.0056 | ** | | | | |
| 6 | A1 | ICSS<br>PPB | GABA | NAc dorsal msh | stim | ICSS<br>PPB | 6 | 2 | 3.848<br>5.978 | 0.5328<br>0.7056 | *t*-test | | t(5)= 5.228 | 0.0034 | ** | | | | |
| | A2 | RTPP 5s<br>Passive 5s | GABA | NAc dorsal msh | stim | RTPP 5s - early<br>Passive 5s - early<br>RTPP 5s - late<br>Passive 5s - late | 6 | 4 | 3.472<br>6.234<br>1.045<br>6.069 | 0.6044<br>0.7374<br>0.2018<br>0.8148 | RM 2-way ANOVA | time period<br>assay type<br>interaction | F (1, 5) = 5.677<br>F (1, 5) = 30.99<br>F (1, 5) = 31.74 | p = 0.0630<br>p = 0.0026<br>p = 0.0024 | ns<br>**<br>** | Sidak | RTPP vs. passive - early<br>RTPP vs. passive - late | 0.0004<br>< 0.0001 | ***<br>**** |
| | B1 | ICSS<br>PPB | GABA | NAc ventral msh | stim | ICSS<br>PPB | 5 | 2 | 5.65<br>10.35 | 0.7<br>1.7 | *t*-test | | t(4)= 3.925 | 0.0172 | * | | | | |
| | B2 | RTPP 5s<br>Passive 5s | GABA | NAc ventral msh | stim | RTPP 5s - early<br>Passive 5s - early<br>RTPP 5s - late<br>Passive 5s - late | 5 | 4 | 5.12<br>9.11<br>1.44<br>8.19 | 0.60<br>1.41<br>0.17<br>1.88 | RM 2-way ANOVA | time period<br>assay type<br>interaction | F (1, 4) = 9.134<br>F (1, 5) = 10.42<br>F (1, 5) = 31.60 | p = 0.0391<br>p = 0.0320<br>p = 0.0049 | *<br>*<br>** | Sidak | RTPP vs. passive - early<br>RTPP vs. passive - late | 0.0007<br>< 0.0001 | ***<br>**** |
| | C1 | Passive 1s | Glut | NAc dorsal msh | stim | pre<br>stim | 4 | 2 | -0.02446<br>-1.984 | 0.2786<br>0.3537 | *t*-test | | t(3)= 3.165 | 0.0253 | * | | | | |
| | | RTPP 1s on | Glut | NAc dorsal msh | stim onset | pre<br>stim | 4 | 2 | 0.5532<br>-0.9017 | 0.5532<br>0.5133 | *t*-test | | t(3)= 1.587 | 0.1054 | ns | | | | |
| | C2 | Passive 5s | Glut | NAc dorsal msh | stim | pre<br>early<br>late | 4 | 3 | -0.2907<br>-1.845<br>-2.014 | 0.3<br>0.3992<br>0.6026 | RM 1-way ANOVA | time period | F(2,3)=7.314 | 0.0246 | * | Holm-Sidak | pre vs early<br>pre vs late<br>early vs late | 0.0403<br>0.0394<br>0.7457 | *<br>*<br>ns |
| | D1 | Passive 1s | Glut | NAc ventral msh | stim | pre<br>stim | 4 | 2 | 0.3371<br>1.813 | 1.813<br>0.4545 | *t*-test | | t(3)= 2.692 | 0.0371 | * | | | | |
| | | RTPP 1s on | Glut | NAc ventral msh | stim onset | pre<br>stim | 4 | 3 | 0.3339<br>2.057 | 0.3344<br>0.2683 | *t*-test | | t(3)= 3.329 | 0.0224 | * | | | | |
| | D2 | Passive 5s | Glut | NAc ventral msh | stim | pre<br>early<br>late | 4 | 3 | -0.2797<br>1.627<br>2.507 | 0.3791<br>0.4065<br>0.3082 | RM 1-way ANOVA | time period | F(2,3)= 20.13 | 0.0022 | ** | Holm-Sidak | pre vs early<br>pre vs late<br>early vs late | 0.0108<br>0.0024<br>0.0979 | *<br>**<br>ns |

**Table S2. Statistics table.** Related to Fig. 1 to 6.
